# Histone H2A serine-1 phosphorylation is a chaperone dependent signal for dimerization with H2B and for enhanced deposition

**DOI:** 10.1101/2022.04.20.488954

**Authors:** Takashi Onikubo, Wei-Lin Wang, David Shechter

**Affiliations:** Department of Biochemistry, Albert Einstein College of Medicine, Bronx, NY 10461

**Author notes:** Contributed equally (Takashi Onikubo, Wei-Lin Wang). CureVac AG, Tübingen, Germany.

## Abstract

Multiple histone chaperones and histone modifications are involved in the folding, transport, and release of histones onto newly replicated DNA. Little is known about histone H2A-H2B pre-deposition histone modifications and their regulation of histone deposition. We previously showed that H2A serine 1 phosphorylation (H2AS1ph) is enriched on the soluble egg histones and on zygotic chromatin in *Xenopus* embryos. Here, we demonstrate that H2AS1 phosphorylation is required for a timely incorporation of H2A-H2B into the pronuclear chromatin. Our analysis revealed that exogenous H2AS1A-H2B dimers were poorly incorporated into pronuclei in egg extract compared with wildtype and H2AS1E-H2B dimers. Chaperone-mediated deposition using histones purified from pronuclei showed that neither Nap1 nor Nucleoplasmin (Npm2) histone deposition was directly affected by endogenous histone posttranslational modification. We further demonstrate that H2AS1 phosphorylation was dependent on Npm2 and required H2B. Surprisingly, Nap1 was incapable of promoting H2AS1 phosphorylation. These results suggest that serine 1 phosphorylation signals a specific state of H2A-H2B dimer bound by Nucleoplasmin. Neither Npm2 nor Nap1 exhibited preference for binding H2AS1A or H2AS1E mutant histones or dimers with H2B *in vitro*. We propose that H2AS1 phosphorylation is a pre-deposition modification that signals for the proper dimerization of H2A-H2B, which in turn activates downstream effectors leading to H2A-H2B deposition.

## INTRODUCTION

In the eukaryotic nucleus 146bp of DNA is wrapped around two of each histones H2A-H2B and H3-H4 to form the fundamental repetitive unit known as a nucleosome (1-3). Before histones are deposited onto DNA, both H2A-H2B and H3-H4 are synthesized and assembled into respective dimers in the cytoplasm (4-6). A detailed analysis of H3-H4 dimers in the cytoplasm has revealed that they are bound by multiple molecular and histone chaperones to ensure the proper assembly of the dimer to be transported into the nucleus, assembled into a tetramer, and then deposited into nucleosomes (4).

The organization of DNA into nucleosomes is critical as it not only allows the long stretch of DNA to be housed inside the nucleus but as it also serves as the fundamental regulatory center for gene expression (2, 7, 8). Consequently, the expression of histones is also a tightly regulated process, in which the canonical histones are expressed only during S-phase (9-11). To achieve this strict regulation, histones interact with various histone chaperones and utilize histone modifications. A cascade of chaperones, such as NASPs, Asf1 and CAF-1, ensures the dimerization of H3-H4 and proper timing for the nuclear import (4, 12, 13). We showed that histone chaperones are themselves regulated on their intrinsically disordered regions (14, 15). Furthermore, acetylation of lysines 5 and 12 by HAT1 has been shown to facilitate the transfer of H3-H4 to Asf1, which is the chaperone directly responsible for the nuclear transport (4, 12, 16-18).

Nap1 has been demonstrated to be the histone chaperone responsible for the nuclear transport of histones H2A-H2B in budding yeast (5, 6) and has been shown to deposit H2A-H2B onto DNA *in vitro* (19-22). However, it still remains unclear if Nap1 is directly responsible for H2A-H2B deposition onto chromatin and if a similar chaperone and modification network exists for H2A-H2B dimer assembly and quality control (23).

During *Xenopus* development, transcription is shut off in the cells until the mid-blastula transition. During the early stages of development, cells rely on the maternally deposited proteins and mRNA to support their accelerated replication (24-26). Histones are also stored in a copious amount in the egg and are constantly replenished through translation (25, 27). We previously reported that serine 1 (S1) phosphorylation is enriched on the soluble pool of H2A in the egg and in the embryonic chromatin (27). Here we report that S1 mutation to alanine (A) significantly hinders the incorporation of exogenous H2A into the pronuclear chromatin formed using *Xenopus* egg extract, despite that purified pronuclear histones, containing S1 phosphorylated H2A, do not enhance *in vitro* nucleosome assembly by Nap1 or Nucleoplasmin (Npm2; rNpm is recombinant Npm2 and eNpm is Npm2 from the egg), H2A-H2B histone chaperones present in the egg. To characterize the conditions necessary for S1 phosphorylation, we employed biochemical analyses using *Xenopus* egg extract to further demonstrate that H2A S1 phosphorylation is dependent on Npm2 and on the presence of H2B, despite Npm2’s ability to bind H2A alone. We also observed that Nap1 was incapable of promoting S1 phosphorylation, despite its presence in the egg. We propose that H2A S1 phosphorylation is a pre-deposition histone modification that signals the proper assembly of H2A-H2B dimers on Npm2 and activates the downstream events leading to deposition of H2A-H2B onto the newly replicated DNA.

### EXPERIMENTAL PROCEDURES

HSS and LSS Xenopus egg extracts were prepared as previously described (28, 29).

#### Cloning of tagged and mutant H2As

A StrepII_tag was inserted inframe in the Cterminus of pRUTH5-*Xl*H2A (containing an N-terminal His6-tag and TEV cleavage site) with the SLiCE cloning method (30). The same cloning method was used to modify pRUTH5_Xl_H2A_StrepII into the Ser-to-Ala mutant, pRUTH5_Xl_H2A_S1A_StrepII, and Ser1-to-Glu phosphomimetic mutation, pRUTH5_Xl_H2A_S1E_StrepII.

#### Protein Purification

Npm2 was purified from laid eggs as previously described (31). Briefly, the crude egg extract was heated at 70°C, and the supernatant was sequentially loaded onto DEAE and phenyl-sepharose columns. Purified egg Npm2 (eNpm) was stored in a buffer (10mM Tris pH8.0, 1mM EDTA, 100mM NaCl, and 1mM DTT). Recombinant Npm2 (rNpm) contained 6XHis tag and purified with Ni-NTA resin (Pierce). The tag was cleaved with TEV protease and further purified over MonoS column (GE) as previously described (31) and stored in the same buffer.

#### X. laevis

Nap1 was cloned from pET51b-Nap1 (a kind gift of R. Heald) into pRUTH5 vector, containing a 6XHis tag followed by a TEV digestion site (31). *Xl* Nap1 was expressed in Rosetta 2 cells (Novagen) for three hours at 37°C. The cells were lysed in (25mM Tris pH8.0, 500mM NaCl, 5mM imidazole, 5mM β-mercaptoethanol, and 1mM PMSF), and the cleared lysate was loaded onto Ni-NTA resin (Pierce). The protein was eluted with a buffer (20mM Tris pH8.0, 200mM NaCl, and 5mM β-mercaptoethanol) containing a stepwise concentration gradient of imidazole between 25-300mM. For the removal of 6XHis tag, EDTA and DTT were added to the final concentrations of 1mM and 2mM, respectively, and the purified Nap1 was treated with TEV protease 1/100 the mass of the purified Nap1 for two days at 4°C. The TEV digested Nap1 was buffer exchanged to the lysis buffer using Amicon Ultra-15 (Millipore) protein concentrator with 10kDa MWCO. The TEV digested protein sample was reloaded onto Ni-NTA to remove undigested protein, and the flowthrough fraction was collected. The protein sample was then buffer exchanged to (10mM Tris pH8.0, 1mM EDTA, 100mM NaCl, and 1mM DTT). The protein sample was then loaded onto Mono Q column (GE) and eluted between the NaCl concentrations of 100mM and 750mM. The fractions containing Nap1 was collected, concentrated, and buffer exchanged to (20mM Tris pH8.0, 100mM NaCl, 1mM EDTA, and 1mM DTT). The concentration of Nap1 was determined using A_280_.

Recombinant histones H2A and H2B were purified as previously described (31). Briefly, all H2A proteins were expressed in BL21(DE3)Star cells (Invitrogen) and inclusion bodies were first purified with Ni-NTA resin and eluted in D1000 buffer (50mM Tris pH8, 1M NaCl, 10 mM betaMercaptoethanol, 6.3M Guanidine-HCl) with 300mM imidazole. DTT was added to 2mM final concentration and digested with TEV protease 1/50 the mass of H2A_StrepII. Uncleaved H2A_StrepII and TEV were removed by subtractive Ni-NTA purification. Equal molar H2A-StrepII and H2B were mixed in HDB2000 buffer and dialyzed against buffers with decreasing salt concentration. H2A-H2B dimers were purified on a Superdex 75 column and concentrated in a 10k MWCO concentrator (Corning). Final protein concentration was measured by UV absorption at 280nm.

#### Pronuclear assembly assay and pronuclear histone purification

pronuclear assembly assay was performed as previously described (27). For pronuclear histone purification, pronuclear assembly was performed with 20ml of LSS divided into 10 tubes with 4,000 demembranated sperms/μl of LSS. After two hours of incubation, the pronuclei were lysed with ELB-CIB (10mM HEPES pH7.8, 50mM KCL, 2.5mM MgCl_2_, 0.25M sucrose, 1mM DTT, 0.1% Triton X-100) supplemented with 1X HALT phosphatase inhibitor cocktail, 1X protease inhibitor cocktail, and 10mM sodium butyrate without spermine and spermidine (29). The pronuclear chromatin was collected by centrifugation over ELB-CIB with 50% glycerol. The chromatin pellet was washed once in ELB-CIB with 300mM KCl and once with a medium salt buffer (20mM HEPES pH7.5, 400mM NaCl, 1mM EDTA, 5% glycerol, and 1mM PMSF) (31). Then the pronuclear chromatin was resuspended in a high salt buffer (20mM HEPES pH7.5, 650mM NaCl, 1Mm EDTA, 340mM sucrose, 5mM β-mercaptoethanol, and 1mM PMSF) and homogenized with a Dounce homogenizer. Homogenized chromatin was dialyzed into HAP low salt buffer (20mM sodium phosphate pH6.8, 600mM NaCl, 5μM β-mercaptoethanol, and 1mM PMSF) and loaded onto hydroxyapatite column (Bio-rad). The full octamer was eluted in a single elution with HAP with 2.5M NaCl, concentrated, and buffer exchanged to (20mM Tris pH8.0, 2M NaCl, 1mM EDTA, 5% glycerol, 10mM β-mercaptoethanol). The concentration of histones was estimated using A280 with calculated molar extinction coefficients of Xenopus core histones.

#### Chromatin assembly assayin vitro

chromatin assembly assays were performed as previously described (31, 32). For chromatin assembly assay, the efficiency is severely affected by the histone:DNA mass ratio (32). Therefore, the mass of pronuclear histones for optimal chromatin assembly was empirically determined by performing a chromatin assembly assay using 20:1 mass ratio of recombinant *Xenopus* Nap1 to plasmid DNA (pGIE0) and titrating amounts of pronuclear histones (see main text). The optimal mass ratio was determined to be 1.4:1 pronuclear histones:DNA, and this mass ratio was used for further chromatin assembly assays.

For the comparative chromatin assembly assay between HeLa core histones and pronuclear histones, Nap1 was included with 10:1 mass ratio with respect to HeLa core histones for both reactions with HeLa core histones and with pronuclear histones. Although 1.4:1 histone:DNA mass ratio was used for pronuclear histones, it was denoted 10:1 Nap1:histone mass ratio for simplicity.

#### In vitro phosphorylation of H2A

5μM of recombinant H2A, H2B, and chaperone at indicated concentrations were mixed in a buffer (10mM HEPES pH7.8, 50mM KCL, 2.5mM MgCl_2_, 1mM DTT) with 1X HALT phosphatase inhibitor cocktail (Pierce) and 1X ATP regeneration system (10mM creatine phosphate (Roche), 10μg/ml creatine kinase (Roche), 2mM ATP (Sigma), 2mM MgCl_2_, 5mM HEPES pH7.5, and 1mM DTT). Then either 2μl or 1μl of egg extract (HSS) was added per 20μl reaction and incubated at 22°C. The reaction was terminated by taking the sample into 1X laemmli sample buffer and snap freezing in liquid nitrogen.

#### Co-precipitation assay of histones

for *in vitro* binding assays, 0.5μM each of H2A-SII and H2B, H2A-SII alone, or untagged H2A and H2B (as non-specific binding control) were mixed with an equimolar amount of indicated chaperone in a buffer (20mM Tris pH7.5, 150mM NaCl, 1mM EDTA, 0.01% NP-40, 10μg/ml BSA, and 1mM DTT). The complex was precipitated with 50μl of Streptactin resin slurry, washed twice in a buffer, and eluted by boiling in 1X laemmli buffer. For binding analysis during S1 phosphorylation, S1 phosphorylation was set up as described earlier with 5μM of either H2A-SII and H2B or untagged H2A and H2B (as non-specific binding control) and in the buffer supplemented with 10μg/ml BSA. The precipitation was performed as for *in vitro* binding assays.

#### Histone peptide pulldown

Biotin-labeled peptides were immobilized on Dynabeads M280 in 1XPBS for 2hr at RT and washed with 1XPBS containing 0.1% TritonX-100 to remove unbound peptides. eNpm and rNap1 were preincubated in b 150 mM KCl, 0.1% TritonX-100 and 10 mM Neutravidin to prevent non-specific binding. Pre-treated eNpm was then incubated with peptide-bound beads in binding buffer (150 mM KCl, 0.1% TritonX-100, 1mM DTT) for one hour. This was followed with the addition of pretreated rNap1 to compete for the binding to peptides for another hour. Finally, beads were washed with binding buffer containing 300 mM KCl to remove unbound protein and boiled with sample buffer for Western blot analysis.

#### Chemicals and antibodies

chemicals and reagents were acquired from Sigma, RPI, or Fisher Scientific. The antibodies used in this study include: monoclonal Npm antibody (33), polyclonal rabbit antibody generated against full-length recombinant Nap1, and S1 phosphorylation antibody(34).

#### Images and adjustment

Gel images were obtained using digital tools (Epson V700 scanner and GE LAS-4000) with 16-bit dynamic range. Complete images were levels adjusted as a whole to improve clarity without obscuring, eliminating, or misrepresenting any information present in the original image acquired.

## RESULTS

### H2AS1A mutation attenuates histone deposition

We previously reported that S1 phosphorylation of H2A is enriched on the soluble pool of H2A stored in *Xenopus* eggs and in the chromatin of the embryos. To test the effect of S1 phosphorylation on chromatin assembly in the egg, we performed a pronuclear assembly assay using low-speed extract of *Xenopus* eggs (LSS) supplemented with preassembled H2A-H2B dimers harboring either S1A or S1E mutation and StrepII tag (“SII”) on the C-terminus of H2A (**Figure 1A**). Collecting the resultant pronuclei and analyzing the chromatin-bound proteins, we confirmed that the wildtype exogenous H2A-SII-H2B dimers were readily incorporated into the pronucleus. We observed that PCNA and B4 deposition occurred at a comparable level to the control, indicating proper chromatin formation and DNA replication in the presence of the exogenous H2A-SII-H2B dimers. We also confirmed that S1 phosphorylation of H2A occurred readily with the exogenous H2A-SII-H2B dimers (**Figure 1B**). However, using the H2A S1A mutant, we observed that histone deposition was reduced compared to wildtype H2A (**Figure 1A and 1C**). PCNA and B4 were deposited onto the chromatin but at a significantly lower level. This slower H2A deposition was relieved by the S1E phospho-mimetic mutation (**Figure 1A**), indicating that S1 phosphorylation is important for timely chromatin assembly in egg extract.

**Figure 1.**
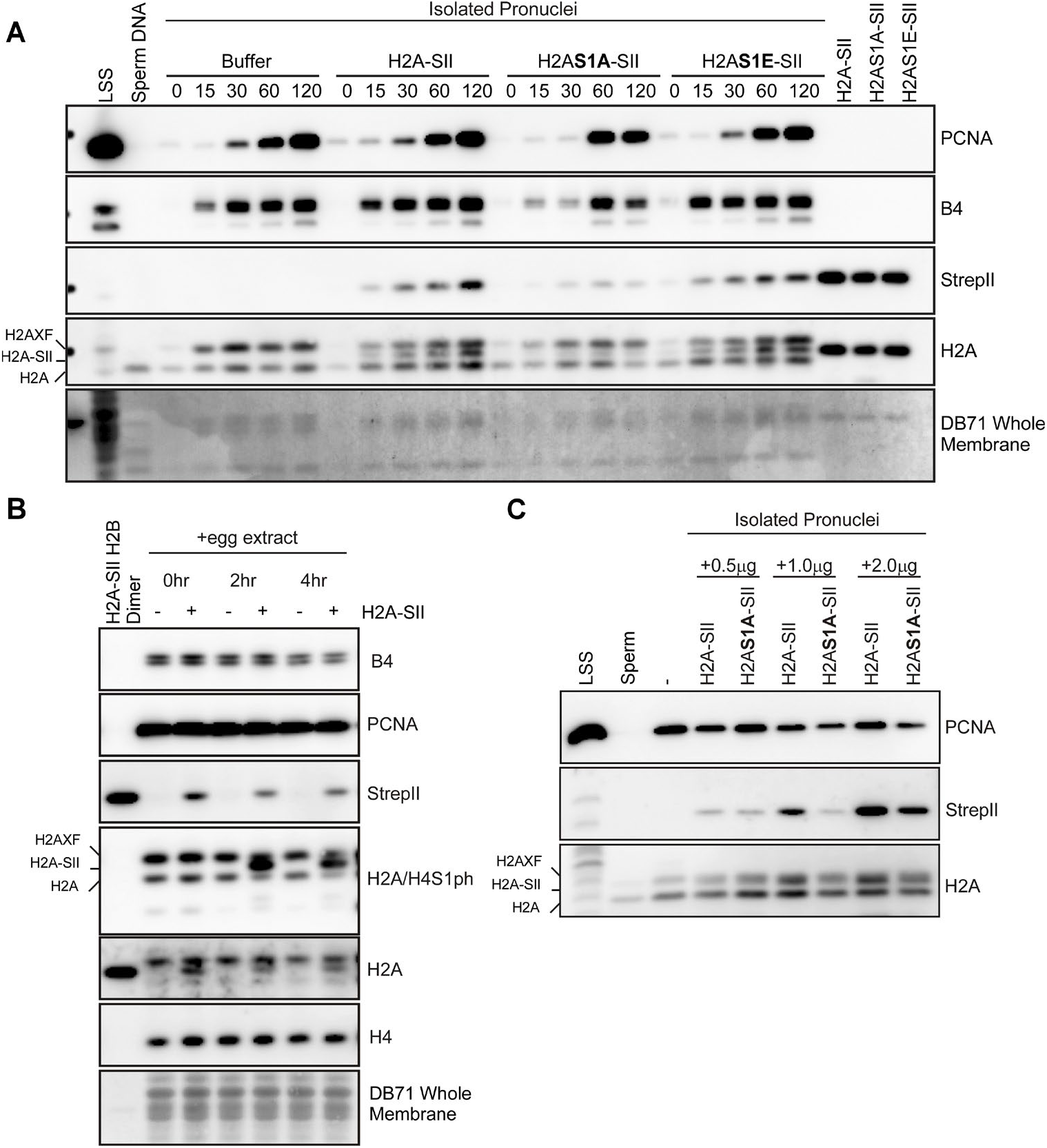
H2A S1A mutation hindered incorporation of H2A into pronuclear chromatin. **A**. S1 phosphorylation and proper chromatin formation are confirmed in the presence of exogenous H2A-SII. A pronuclear assembly assay is performed with LSS supplemented with H2A-SII-H2B. The isolated pronuclei were probed for B4, PCNA, H2A-SII (Strep-tag II), H2A/H4 S1 phosphorylation, H2A and H4. **B**. Pronuclear assembly assay was performed using LSS supplemented with H2A-SII-H2B, H2AS1A-SII-H2B, H2AS1E-SII-H2B, or buffer. The pronuclei were collected and probed for PCNA, B4, exogenous H2A (Strep-tag II), and H2A by immunoblot. The incorporation of exogenous H2A was slower with H2AS1A mutant compared to wt or H2AS1E. For the anti-H2A blot, the positions of the canonical H2A, H2A-SII, and H2A.X-F are indicated. A whole membrane stain is also shown.

To test if H2A S1 phosphorylation directly enhances the histone deposition, we purified core histones from the pronuclei assembled in extract and tested if the pronuclear core histones would exhibit accelerated histone deposition *in vitro* by chaperones present in the egg. The purified core histones contained a significant level of H2A.X.3 (H2A.X-F) and carried S1 phosphorylation on H2A, H2A.X.3, and H4, as previously reported (**Figure 2A,B**) (27, 28). This observation indicated that the purified pronuclear core histones retained the composition and modifications as in the egg.

**Figure 2.**
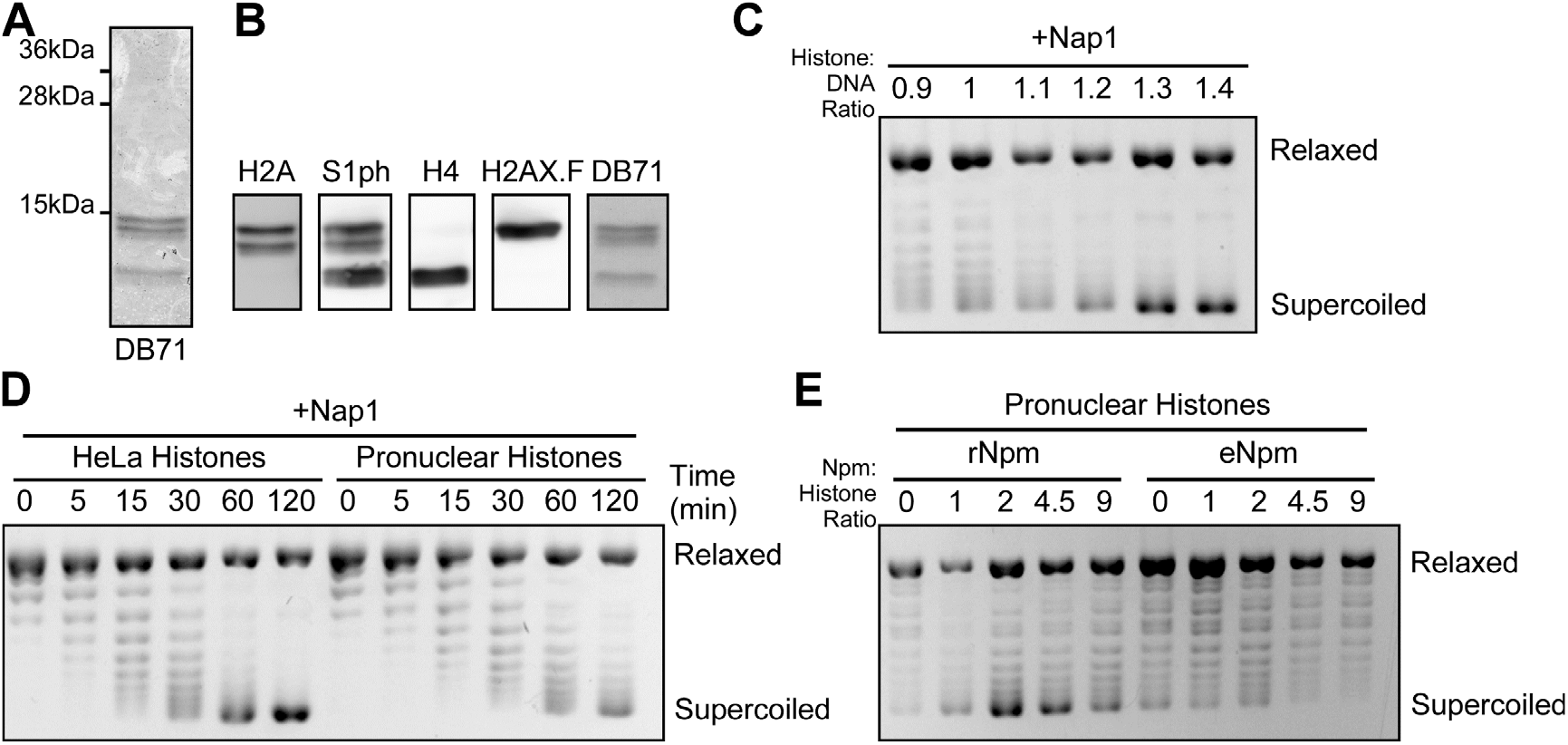
Pronuclear histones with H2A S1 phosphorylation does not promote in vitro histone deposition. *Pronuclear* histones were purified from pronuclei assembled in LSS, and in vitro chromatin assembly assay was performed. A. The purified histones were resolved on SDS-PAGE gel and stained with Coomassie blue. B. Purified pronuclear histones were probed for H2A, S1 phosphorylation, H4, H2A.X-F. Note that H2A antibody showed both H2A and H2A.X-F. The three bands seen with S1 phosphorylation antibody are S1 phosphorylation on H2A.X-F, H2A, and H4. The whole membrane stain is also shown. C. The proper amount of purified pronuclear histones was determined with a chromatin assembly assay using a saturating amount of Nap1 and titrating amounts of pronuclear histones. The optimal amount of pronuclear histones (1.4:1 histone to DNA mass ratio) was used in subsequent chromatin assembly assays. D. Pronuclear histones showed lower efficiency in chromatin assembly with Nap1 compared to HeLa core histones. Chromatin assembly assay was performed using 10:1 mass ratio of Nap1 to histone ratio and with either HeLa core histones or pronuclear histones. The degree of histone deposition, as supercoiling of the plasmid, was measured at indicated time points. E. Pronuclear histones did not reverse the sequestration by eNpm. Chromatin assembly assay was performed with pronuclear histones and titrating amounts of rNpm or eNpm. The amount of Npm is indicated as Npm to histone mass ratios.

For *in vitro* chromatin assays, it is critical to maintain the histone:DNA mass ratio of 1:1 for optimal histone deposition. We empirically determined the optimal pronuclear histone:DNA mass ratio by titrating the pronuclear histones while keeping the constant concentration of DNA and the saturating concentration of *Xenopus* recombinant Nap1, at 20:1 mass ratio with DNA. We observed that pronuclear histones required higher mass ratio, at 1.4:1, compared to 1:1 of HeLa core histones, for the optimal histone deposition (**Figure 2C**). We then reduced the Nap1:histone mass ratio to 10:1, and compared the histone deposition with pronuclear histones to HeLa core histones *in vitro* (**Figure 2D**). We observed a mildly reduced histone deposition rate with the pronuclear histones. Both pronuclear histone and HeLa core histone samples contain an equal amount of Nap1.

Npm2 has been proposed to be responsible for storing histones H2A and H2B in Xenopus eggs (35, 36). We also reported that Npm2 purified from the egg exhibits histone sequestration in a hyperphosphorylation-dependent manner (31). Therefore, we tested to see if the pronuclear histones would reverse the sequestration of egg Npm (eNpm). To address this question, we performed an *in vitro* chromatin assembly assay with purified eNpm and pronuclear histones (**Figure 2E**). However, we still observed strong sequestration by eNpm and mass-ratio-specific histone deposition by recombinant Npm2 as we observed with HeLa core histones (31). These results indicated that pronuclear histones, containing S1 phosphorylation on H2A do not directly affect the histone deposition or sequestration by Nap1 or Npm2.

### S1 phosphorylation of H2A is Nucleoplasmin dependent

To further analyze H2A S1 phosphorylation, we set up an *in vitro* H2A S1 phosphorylation assay using high speed *Xenopus* egg extract (HSS). Since *in vitro* dimerization of H2A and H2B requires a high concentration of NaCl and may interfere with the assay, 5μM of H2A and of H2B were individually mixed with 5μM eNpm followed by addition of 2μl clarified egg extract in a 20μl final reaction volume. Under this experimental setup, we observed strong phosphorylation on S1 of the exogenous H2A after 2 hours (**Figure 3A**).

**Figure 3.**
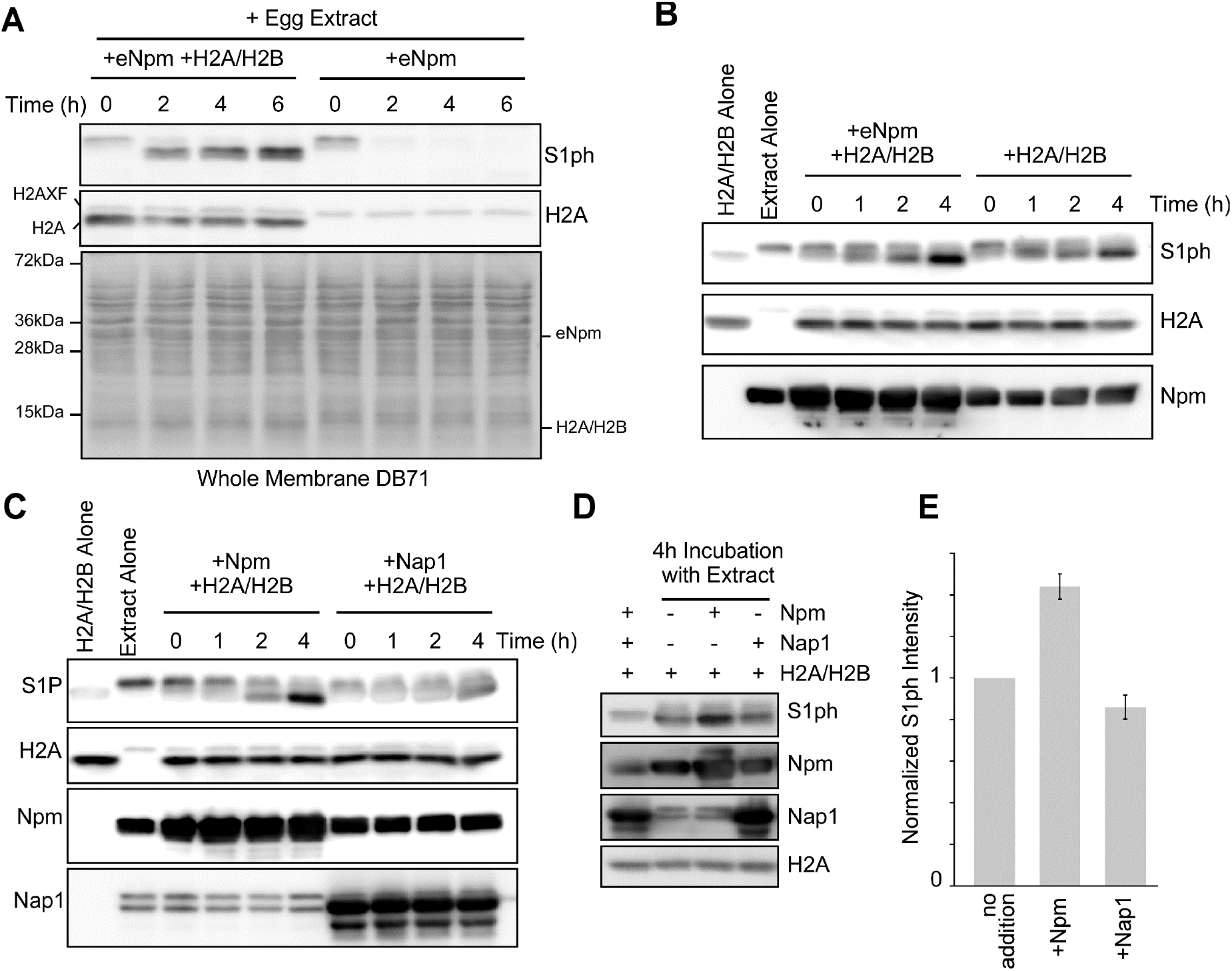
H2A S1 phosphorylation is Npm-dependent. *In vitro* S1 phosphorylation assay was performed using recombinant H2A-H2B and diluted egg extract. **A**. S1 phosphorylation occurred on exogenous H2A. *In vitro* S1 phosphorylation assay was performed with eNpm+H2A/H2B and with eNpm alone. S1 phosphorylation was confirmed by immunoblot using anti-S1P antibody. Note that anti-H2A and anti-S1P antibody recognized H2A.X-F in the extract. The whole membrane stained with DB71 is also shown. **B**. S1 phosphorylation occurred more strongly when H2A and H2B were phosphorylated in the presence of eNpm. S1 phosphorylation assay was performed with H2A/H2B with or without eNpm. Immunoblot for S1 phosphorylation, H2A, and Npm are shown. **C**. Nap1 did not promote S1 phosphorylation. S1 phosphorylation was performed with H2A/H2B supplemented with either eNpm or Nap1. Immunoblot for S1 phosphorylation, H2A, Npm, and Nap1 are shown. **D**. S1 phosphorylation assay was repeated with eNpm, Nap1 and no chaperone. Four hour incubation products were resolved side-by-side on a SDS-PAGE gel. **E**. Quantification of D is shown. The data represents the mean ± standard error of three replicates.

To study the effect of histone chaperones on H2A S1 phosphorylation, we repeated the assay in the presence and absence of eNpm (Note that extract contains a large amount of Npm2, estimated at 0.7mg/ml, *data not shown*). Therefore, the amount of extract was reduced to half for the subsequent assays to minimize the effect of endogenous Npm from the extract.). We observed higher S1 phosphorylation in the presence of exogenous eNpm (**Figure 3B**). We also tested if Nap1 could similarly promote S1 phosphorylation by adding recombinant *Xenopus* Nap1. However, we did not observe any enhancement of S1 phosphorylation compared to no chaperone (**Figure 3C**). Since we observed some S1 phosphorylation in the absence of Npm or in the presence of Nap1, which is likely due to endogenous Npm present in HSS, we initially attempted to perform an S1 phosphorylation assay using Npmand Nap1-depleted HSS. However, the kinase activity was lost during the depletion, even for the mock depletion using resin alone (*data not shown*). Therefore, we repeated the assay and quantified the levels of S1 phosphorylation in the anti-S1P immunoblot. Comparing the samples with eNpm, Nap1, and without exogenous chaperones we confirmed a significant increase in S1 phosphorylation with eNpm and a mild reduction with Nap1 (**Figure 3D and E**). These results strongly suggested that S1 phosphorylation is Npm2-dependent.

### S1 phosphorylation signals for the dimerization of H2A-H2B on Nucleoplasmin

To delineate the function of S1 phosphorylation, we probed further conditions potentially required for S1 phosphorylation. Since Npm2 serves as a storage histone chaperone in the egg, we hypothesized that S1 phosphorylation may signal for the proper dimer assembly on Npm2. To test this hypothesis, we compared S1 phosphorylation in the sample containing both H2A and H2B to the sample containing H2A alone. Consistent with our hypothesis, the absence of H2B reduced *de novo* S1 phosphorylation on exogenous H2A (**Figure 4A**). This result suggested that S1 phosphorylation is dependent on Npm binding both H2A and H2B.

**Figure 4.**
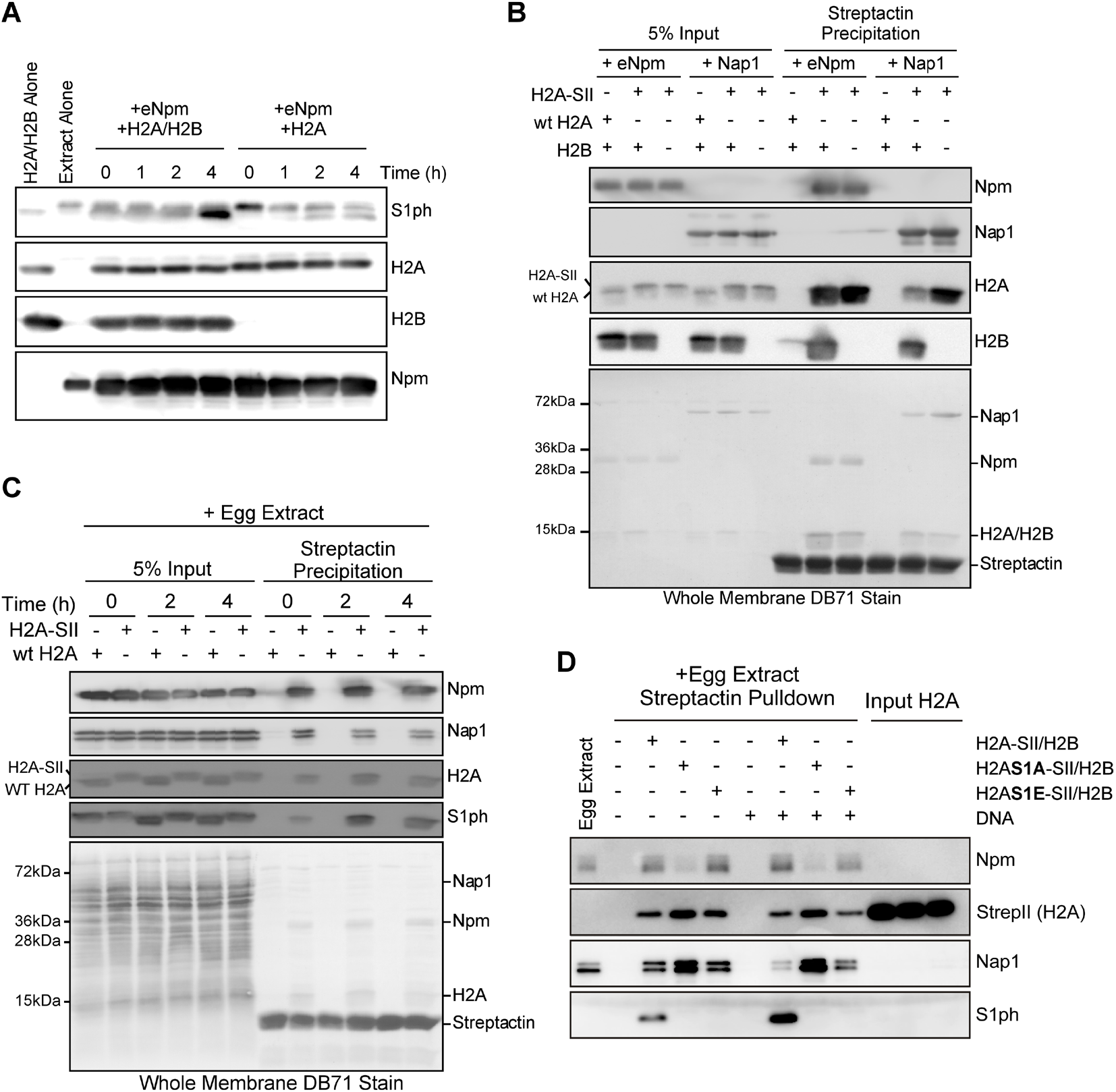
H2A S1 phosphorylation is co-dependent on H2B. **A**. S1 phosphorylation was performed with either H2A/H2B or H2A alone supplemented with eNpm and probed at indicated time points. Immunoblot for S1 phosphorylation, H2A, H2B and Npm are shown. **B**. Both eNpm and Nap1 bound tightly to H2A-SII/H2B and to H2A-SII alone *in vitro*. eNpm and Nap1 were mixed with either H2A-SII alone or with H2A-SII and H2B. The complex was precipitated with Streptactin, and the association with respective chaperones was confirmed by immunoblot. The samples containing tag-less wildtype H2A served as non-specific interaction control. Immunoblot for Npm, H2A, H2B, and the whole membrane stain are shown. **C**. Npm and Nap1 interact with H2A during S1 phosphorylation. S1 phosphorylation was performed with H2A-SII/H2B and without any exogenous chaperone. The chaperone-H2A-SII/H2B complex was precipitated with Streptactin at indicated time points and was probed for Npm and Nap1. Immunoblot for H2A, S1 phosphorylation, and the whole membrane stain are shown. **D**. StrepII-tagged H2A/H2B dimers, with wildtype or S1A or S1E mutant H2A, were incubated in egg extract in the absence or presence of DNA. The chaperone-H2A-SII/H2B complex was precipitated with Streptactin and immunoblotted for Npm, StrepII tag, Nap1, and S1ph as indicated.

It was also possible that Npm is incapable of binding H2A alone, and the loss of signal with H2A alone may have been a result of this absence of interaction. To test this possibility, we performed a precipitation assay using recombinant H2A labeled with StrepII tag on the C-terminus mixed with either eNpm or Nap1 *in vitro*, while the sample with unlabeled H2A served as a negative control (**Figure 4B**). We confirmed that both Npm and Nap1 efficiently co-precipitated with both H2A alone and H2A-H2B dimers, indicating interaction with both chaperones with H2A alone. To test if H2A-H2B is preferentially bound by either Npm or Nap1 during S1 phosphorylation, we also performed a similar precipitation assay over the course of *in vitro* S1 phosphorylation assay in a small quantity of egg extract without any exogenous chaperone (**Figure 4C**). Although some interaction was seen with Nap1, we confirmed a strong interaction between H2A-H2B and Npm at all time points tested (between 0 to 4 hours), while S1 phosphorylation was also observed. Furthermore, we observed that the amounts of co-precipitated Npm and Nap1 remained quite constant at all time points tested. This observation suggested that S1 phosphorylation did not specifically cause the transfer of histones from one histone chaperone to another under this experimental setting using diluted egg extract. This result was consistent with our *in vitro* chromatin assembly assays demonstrating that S1 phosphorylation not directly causing an accelerated histone release from Nap1 or from eNpm (**Figure 2D and E**).

In egg extract H2A phosphorylation may be necessary for it to associate with Npm as Npm precipitation was specifically reduced in a pulldown assay using H2AS1A-dimer and undiluted egg extract, while Npm was strongly precipitated by H2A and H2AS1E (**Figure 4D**). Conversely, Nap1 was enriched with the H2AS1-dimer in undiluted extract, suggesting that another unidentified factor may be promoting the interaction of H2AS1A to Nap1.

### H2A S1ph containing peptides or H2AS1 mutants do not preferentially bind Npm or Nap1 in vitro

To test if H2AS1 phosphorylation has a local effect on interaction with chaperones we performed a number of peptide pulldown interaction studies. First, we used biotinylated H3 (44-63), H2A (1-22), and H2AS1ph (1-22) peptides and incubated them with Npm, Nap1, or both to determine relative affinities (**Figure 5A**). Both Npm and Nap1 bound the N-terminal H2A peptides and H3 peptides as previously reported, although Npm clearly showed stronger interaction with H2A peptide (20, 37). We first observed that both Npm and Nap1 interacted with S1 phosphorylated and unphosphorylated H2A peptides with similar affinities. As we mixed Npm with Nap1, we observed that S1 phosphorylated H2A peptide was more efficiently competed off of Npm by Nap1. This result showed that the N-terminal tail of H2A indeed interacts with Npm and phosphorylation at that site mildly reduces the affinity.

**Figure 5.**
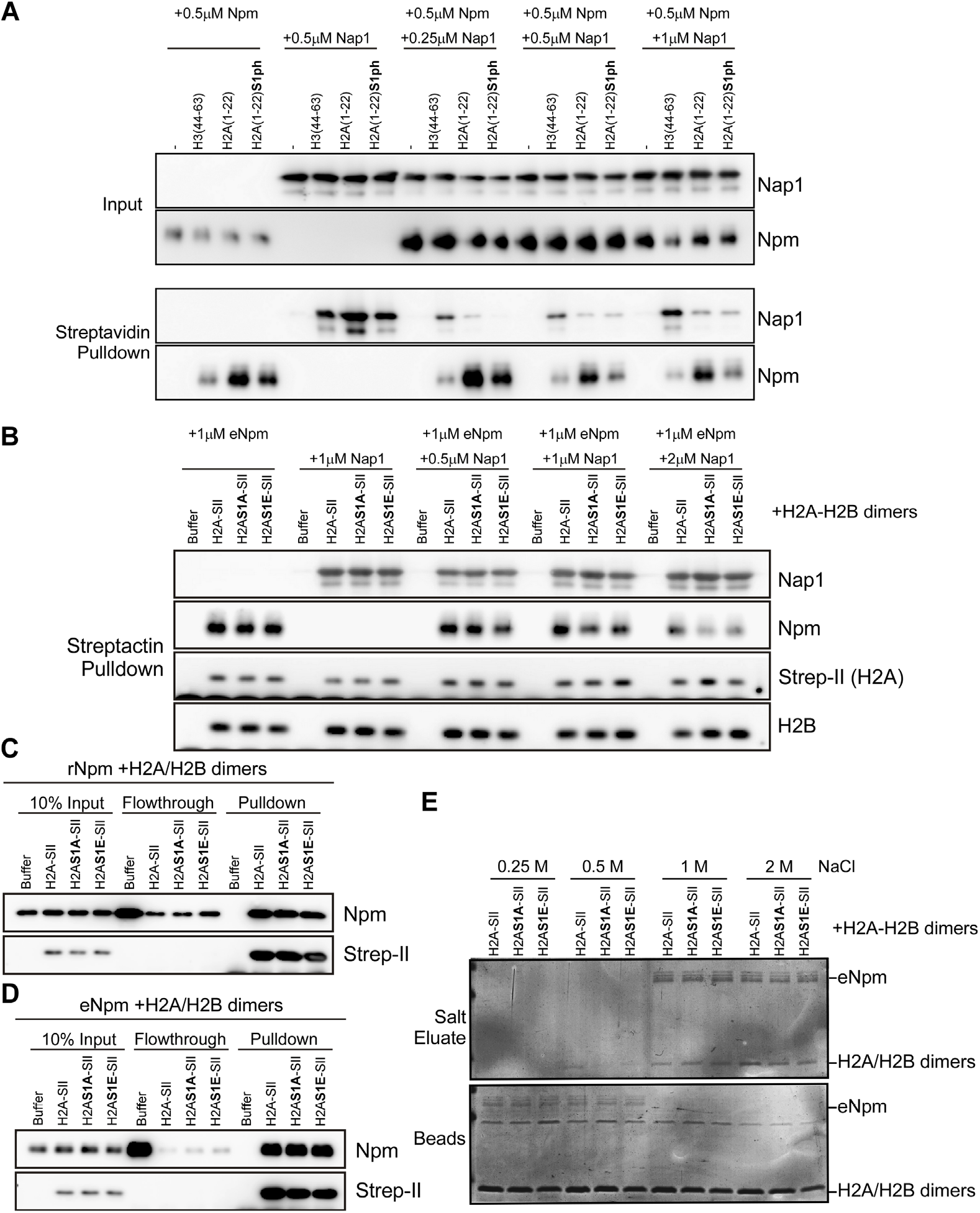
S1 phosphorylation reduces the local affinity of H2A N-terminal tail to Npm. **A**. A competitive pepetide pulldown assay was performed with biotinylated H3 (44-63), H2A (1-22), and H2A (1-22) S1phosphorylated peptides with a constant concentration of Npm and titrating concentrations of Nap1. Pepetide pulldown of Npm and Nap1 alone are also showed. **B**. A competitive pulldown assay was performed using StrepII-tagged full-length H2A-H2B dimers with wildtype H2A (H2A-SII), S1A (H2AS1A-SII), and S1E (H2AS1E-SII). Immunoblot for H2A with Strep-II and H2B are also shown for the efficiency of pulldown and dimerization of H2A-H2B. **C**.and **D**. S1 phosphorylation does not affect the overall affinity for H2A-H2B to rNpm and eNpm. Co-precipitation assay was performed with StrepII-tagged H2A-H2B dimers (wildtype, S1A, and S1E) and rNpm (C.) or eNpm (D.). **E**. The overall affinity of Npm to H2A-H2B is not affected by S1 phosphorylation. The Npm-H2A-H2B complexes with wildtype, S1A, and S1E were immobilized on beads through StrepII tag on H2A. Then Npm was eluted with indicated concentrations of NaCl.

To test if similar effect is observed with fulllength proteins, we performed a competitive pulldown assay with wildtype H2A-H2B dimers or H2AS1A and phosphomimetic S1E mutants tagged with StrepII (**Figure 5B**). Surprisingly, we observed that both S1A and S1E mutants were more efficiently competed off of Npm by Nap1 compared to wildtype. This result suggested that the hydroxyl group of H2A S1 may be promoting the interaction between Npm and H2A N-terminal tail.

Since we previously demonstrated that the post-translational modifications on egg Npm (eNpm) enhanced the affinity for histones (31), we compared eNpm and unmodified rNpm in a pulldown assay with the tagged dimers. We did not observe difference in Npm enriched in these pulldowns (**Figure 5C, D**). Finally, to more directly ascertain relative affinities of eNpm for the H2A-H2B dimers, we bound eNpm to them and eluted with increasing salt concentrations. We did not observe any difference in interaction, as all three dimers bound eNpm at 0.5M NaCl and did not bind at 1M NaCl (**Figure 5E**). These results showed that S1 phosphorylation has a similar effect on the overall affinity of Npm to H2A-H2B regardless of post-translational modifications on Npm.

### Nucleoplasmin and Nap1 are independently dispensable for chromatin assembly in egg extract

To determine if Npm is directly involved in chromatin assembly, we immunodepleted Npm from egg extract (**Figure 6A**) and performed chromatin assembly assay. Npm was quantitatively removed from extract without affecting the levels of Nap1 or histones. Npm-depleted extract supercoiled plasmid DNA identically to the control, and addition of exogenous eNpm did not influence the rate (**Figure 6B**).

**Figure 6.**
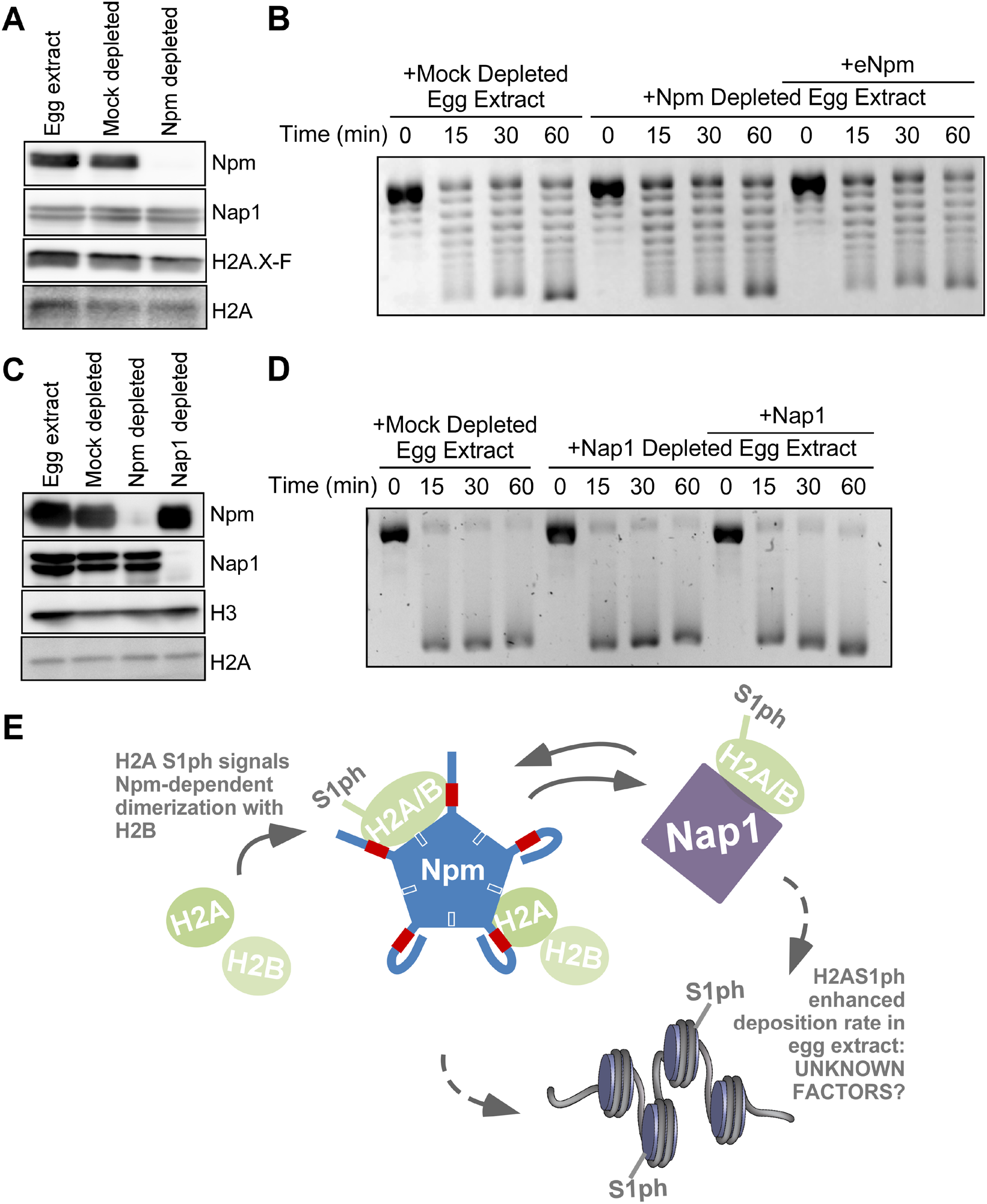
Immunodepletion of Npm and Nap1 from extract did not affect chromatin assembly. **A**. Npm was selectively depleted from extract. An immunoblot for Npm, Nap1, H2A.X-F, and H2A is shown for the untreated, mock depleted, and Npm depleted extracts. **B**. Chromatin assembly assay was performed with mock depleted, Npm depleted extracts, and Npm depleted extract supplemented with purified eNpm. **C**. Nap1 was selectively depleted from extract. Immunoblot for Npm, Nap1, H3 and H2A is shown as in A. **D**. Chromatin assembly was performed with mock depleted, Nap1 depleted extracts, and Nap1 depleted extracts supplemented with purified recombinant Nap1. **E**. Our model for the function of S1 phosphorylation is presented. S1 phosphorylation signals the dimerization of H2A and H2B on Npm and channels the dimer into downstream activities leading to deposition onto DNA.

In parallel, we performed a similar study to test the necessity for Nap1 in chromatin assembly and observed that extract depleted for Nap1 (**Figure 6C**) was also capable of supercoiling plasmid DNA. Re-addition of exogenous Nap1 did not affect this chromatin assembly (**Figure 6D**).

## DISCUSSION

Previously, we reported that H2AS1 phosphorylation accumulates on the soluble pool of H2A in *Xenopus* eggs and in the zygotic chromatin (27). Here we showed that the S1 phosphorylation is dependent on the interaction of H2A with Npm and its dimerization with H2B. We also demonstrated that H2AS1ph is correlated with more rapid deposition into chromatin. Nap1 is the other major chaperone that binds to H2A/H2B in *Xenopus* eggs, yet H2AS1 phosphorylation primarily occurs on Npm. Both chaperones are dispensable for chromatin assembly in cell free egg extract, highlighting a mechanism in which H2A-H2B dimerization is signaled by phosphorylation when bound to Npm, but in equilibrium with Nap1, and the possibility of yet another downstream effector directly responsible for histone deposition (summarized in **Figure 6E**).

### Maternal histone chaperones and deposition

Several histone chaperones have been identified for H2A-H2B, such as Nap1, Npm, and FACT(5, 19, 37-42). Npm is a maternal histone chaperone responsible for the storage of H2A-H2B during early development (31, 43) and the period of transcriptional quiescence prior to zygotic gene activation (25, 26). Therefore, the coordination between timely release and secure storage of histones is crucial for establishing and maintaining the zygotic epigenome. Our observation that S1A mutation reduced the incorporation of H2A into the pronucleus strongly suggests that S1 phosphorylation is a pre-deposition histone modification signal leading to further processing and deposition into the zygotic chromatin, similar to H4 K5 and 12 acetylation observed in the somatic cells.

We also demonstrated that S1 phosphorylation is dependent on H2B and Npm. Along with our observation that Npm readily binds H2A alone, this result suggested that S1 phosphorylation is signaling H2A-H2B dimer formation on Npm. Since Npm serves as the storage chaperone, it is possible that this signaling for dimerization may be important for recruiting downstream effectors necessary for histone deposition and ensuring that only the functional H2A-H2B is channeled into the histone deposition pathway. Our peptide pulldown analysis also showed that H2A C-terminal tail interacts with Npm and S1 phosphorylation reduces the local affinity of H2A tail to Npm, while the overall affinity of Npm to histones remains largely unchanged. These results are consistent with our model that S1 phosphorylation signals for downstream effectors and suggest that the release of H2A Nterminal tail from Npm is critical for that function. Identification of the processes that follow S1 phosphorylation will be particularly interesting, as it will also shed light on the function of S1 phosphorylation in somatic cells, in which S1 phosphorylation was found enriched on the S-phase and mitotic chromatin (44).

We were surprised that S1 phosphorylation was dependent on Npm and not Nap1 in the eggs. Nap1 is also expressed in somatic cells, but Npm expression is limited to the embryos (45, 46). This result may suggest that another chaperone or possibly Nap1 with specific posttranslational modifications promote predeposition S1 phosphorylation in somatic cells.

Together, we propose S1 phosphorylation is a general pre-deposition histone modification that serves as a quality control crucial for the proper H2A-H2B processing and deposition.

## ACKNOWLEDGEMENTS

This work was supported by the National Institutes of Health [R01GM135614 to D.S.], the Alexander and Alexandrine Sinsheimer Foundation Scholar Award [to D.S.], and the The American Cancer Society

– Robbie Sue Mudd Kidney Cancer Research Scholar Grant [124891-RSG-13-396-01 DMC].

## AUTHOR CONTRIBUTIONS

TO and WW equally conceived of and performed experiments, interpreted results, prepared Figures and co-authored the manuscript. DS conceived of experiments and interpreted results and co-authored the manuscript.

